# Human Assembloid Model of Emergent Neurotropic Enteroviruses

**DOI:** 10.1101/2025.11.18.689148

**Authors:** Christine E. Peters, Jimena Andersen, Min-Yin Li, Lauren Varanese, Mayuri Vijay Thete, Se-Jin Yoon, Taylor Pio, Nicholas Thom, Xiaoyu Chen, Wenjie Qiao, Jan E. Carette, Sergiu P. Paşca

**Author notes:** Correspondence should be addressed to S.P.P. and J.E.C. ( &). Contributed equally (Christine E. Peters, Jimena Andersen, Min-Yin Li).

## Abstract

Enteroviruses (EVs) are the leading cause of viral meningitis in children. Recent outbreaks of non-polio EVs, most notably EV-A71 and EV-D68, have been associated with a polio-like paralysis known as acute flaccid myelitis (AFM). The lack of relevant models that mimic the cellular and functional responses of these human-restricted pathogens has hampered the development of effective treatments. We have previously engineered human stem cell-derived assembloids that recapitulate the neuromuscular connections underlying muscle contractions by integrating human spinal cord/hindbrain organoids (hSpO) and human skeletal muscle. Here, we used organoids and assembloids to investigate polio and non-polio EV pathogenesis. Infection of assembloids with poliovirus (PV), EV-D68 and EV-A71 resulted in loss of muscle contraction for all three viruses, which could be prevented by treatment with an antiviral agent. Yet, despite the convergence on neuronal dysfunction, the cellular targets by which each virus acted differed. More specifically, single cell transcriptomic profiling uncovered divergent cell tropisms between the EVs, and live imaging experiments revealed different modes and kinetics of cell damage. Altogether, we describe a multi-cellular model that captures viral pathogenesis in a human and circuit-relevant context.

## Introduction

Enteroviruses (EVs) are the leading cause of viral meningitis in children. EVs can cause other severe central nervous system (CNS) diseases such as poliomyelitis, encephalitis, and a polio-like paralysis named acute flaccid myelitis (AFM) (Pons-Salort et al., 2015). Poliovirus (PV), the etiological agent of poliomyelitis, is now on the verge of eradication following widespread vaccination (Kishore et al., 2024). As a result, most recent EV infections associated with disabling paralysis are due to outbreaks of non-polio enteroviruses, most notably EV-A71 and EV-D68 (Suresh et al., 2018). The study of how EVs cause AFM and the factors that determine their neuropathogenesis has been limited by the lack of a human model system that recapitulates the cell diversity, local circuitry and cell-cell interactions in the spinal cord and periphery.

We previously developed human spinal cord/hindbrain organoids (hSpO) from induced human pluripotent stem (hiPS) cells that recapitulate the cell diversity of the developing human spinal cord/hindbrain (Andersen et al., 2020), including not only motor neurons, but also glutamatergic and GABAergic interneurons, as well as astrocytes and oligodendrocytes. Importantly, by assembling hSpO with organoids containing differentiated human skeletal myoblasts (hSkM), we showed that motor neurons can innervate muscle cells, form functional neuromuscular contacts and trigger muscle contraction.

Here, we leverage the ability of assembloids to capture complex cell-cell interactions to create models of infection of the neuromuscular unit to study the paralytic enteroviruses PV, EV-D68 and EV-A71. We show that while infection with each of the three EVs results in muscle paralysis, the mode by which each virus exerted this effect differs. Interestingly, we uncovered differences in both cell tropism and cell death kinetics upon infection that ultimately converged on neuronal dysfunction. Altogether, our study provides insight into how EVs cause neuropathogenesis in the CNS and establishes a human platform that could be applied to identify therapeutics for viral infections.

## Results

### *In vitro* models of enterovirus infection in hSpO-hSkM assembloids and spinal cord organoids

We first generated hSpO-hSkM assembloids from hiPS cells and human skeletal myoblasts to probe the effect of EVs on functional contractions (n = 4 hiPS cell lines in various experiments as summarized in **Table S1**). For this, hSpO and hSkM were separately generated from hiPS cells or human skeletal myoblasts, which were directed to differentiate into myotubes within an extracellular matrix. Individual hSpO and hSkM were assembled and hSpO-hSkM allowed to integrate functionally for approximately 2 weeks. At this point, which we labeled as day 0 (d0), we performed live imaging to capture baseline spontaneous contractions. hSpO-hSkM assembloids were then exposed to either PV-1 at 10^4^ PFU, enterovirus A71 (EV-A71) at 10^4^ PFU, or enterovirus D68 (EV-D68) 2014 outbreak strain US/IL/14-18952 at 10^5^ PFU (**Fig. 1A**). Uninfected hSpO-hSkM were used as controls. We confirmed that hSpO-hSkM assembloids were susceptible to EV infection by immunostaining for viral antigens after two days (**Fig. 1B**). Viral antigens were detected primarily in the hSpO region. We monitored contractions in hSpO-hSkM every two days for a total of 8 days and quantified displacement of pixels over time in imaging fields (subdivided into 4 subfields, **Fig. 1C**), as we have previously done (Andersen et al., 2020). We observed a strong reduction in muscle contraction over the course of a week for all three EVs and across multiple hiPS lines (**Fig. 1D–J, Supplemental Videos 1–4**), despite spontaneous activity remaining mostly consistent in uninfected assembloids. Treatment with the antiviral rupintrivir– a viral 3C protease inhibitor with broad anti-picornavirus activity (Binford et al., 2005; Kim et al., 2012; Sun et al., 2015), prevented the loss of contractions, indicating that this phenotype was a consequence of virus infection.

**Figure 1:**
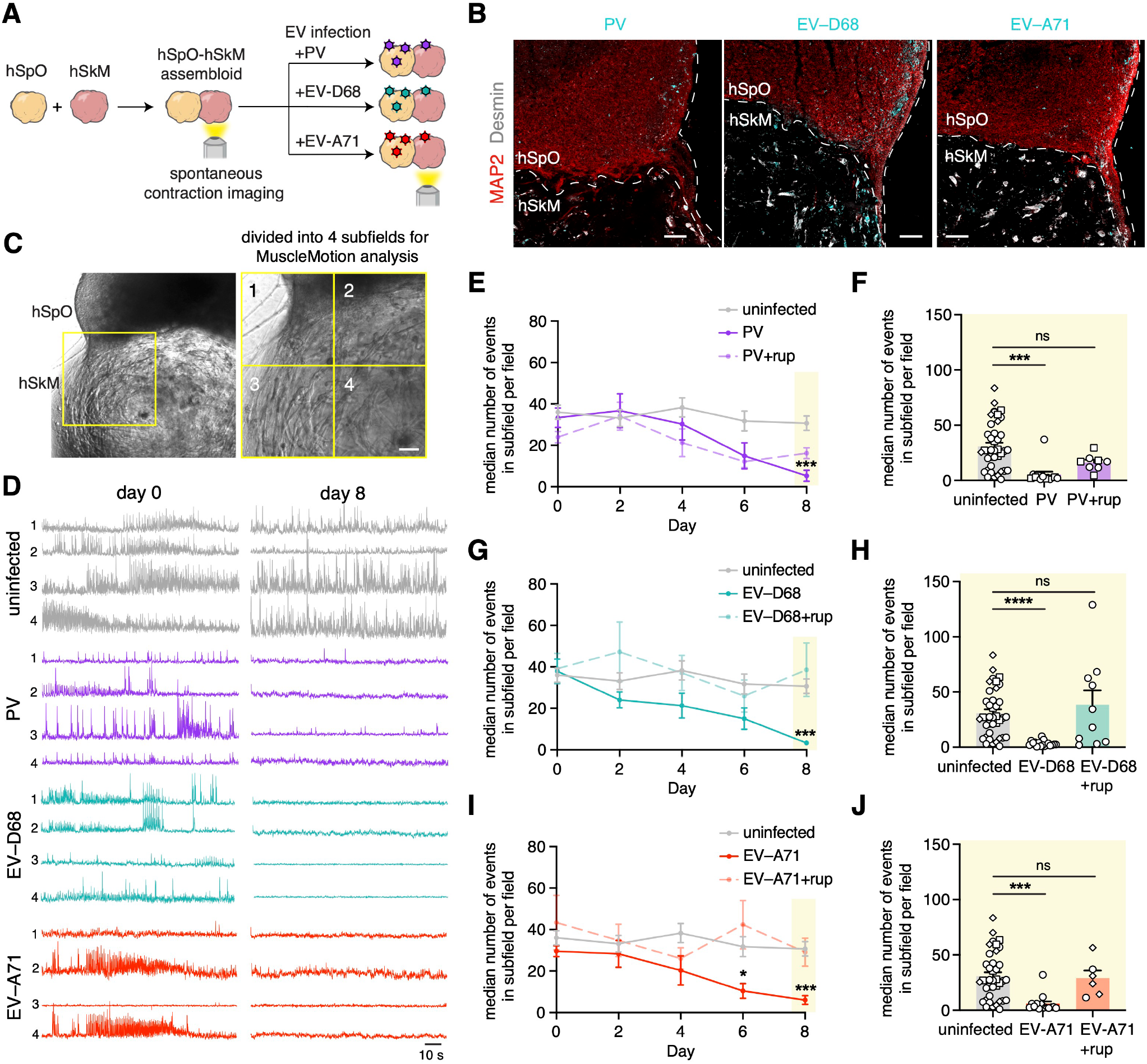
Model of enterovirus-related acute flaccid myelitis using hSpO-hSkM assembloids. **(A)** Schematic showing hSpO-hSkM assembloid setup and infection with polio and non-polio EVs. **(B)** Representative immunofluorescent images of hSpO at two days post infection with PV, EV-D68, or EV-A71. PV, EV-D68 and EV-A71 expression was detected by an anti-dsRNA antibody (cyan). Scale bars, 100 μm. **(C)** Representative bright-field image showing intact hSpO-hSkM assembloid and imaging field for spontaneous muscle contraction analysis. Imaging fields are divided into 4 subfields for analysis. Scale bar, 100 μm. **(D)** Representative spontaneous contraction traces in a subfield of uninfected or infected hSpO-hSkM assembloids. See videos S1-4 for imaging of spontaneous contractions of uninfected or infected hSpO-hSkM at d0, d2 and d8. **(E-J)** Quantification of spontaneous contractions over a 2-min period showing the median number of events in subfields per field (n = 40 fields from 20 assembloids for uninfected hSpO-hSkM from 2 differentiations, n = 13 fields from 7 assembloids for PV infected hSpO-hSkM from 2 differentiations, n = 8 fields from 4 assembloids for PV+rup infected hSpO-hSkM from 1 differentiation, n = 17 fields from 9 assembloids for EV-D68 infected hSpO-hSkM from 2 differentiations, n = 10 fields from 5 assembloids for EV-D68+rup infected hSpO-hSkM from 2 differentiations, n = 14 fields from 8 assembloids for EV-A71 infected hSpO-hSkM from 2 differentiations, n = 6 fields from 3 assembloids for EV-A71+rup infected hSpO-hSkM from 2 differentiations. Individual values from day 8 are graphed in F, H, and J. Datasets represent mean ± SEM. P-values were determined by one-way ANOVA with Holm-Sidak correction. *P<0.05, ***P<0.001, ****P<0.0001. ns, non-significant.

We next performed a thorough characterization of the dynamics of viral infection in hSpO to better understand how divergent EVs cause paralysis. To further investigate the susceptibility of hSpO to infection by EVs, we performed a time-course of infection over ten days. Samples were harvested at 1, 2, 4, 6, 8 and 10 days post-infection (dpi) and analyzed by immunostaining using antibodies against EV capsid proteins or dsRNA. Cells positive for PV, EV-D68 or EV-A71 could be detected as early as 1 dpi, with viral antigens seen in both the cell body and processes (**Fig. 2A**). We assessed viral replication by measuring intracellular viral RNA levels over time. Quantitative RT-PCR of hSpO lysates revealed an increase in viral RNA over time (**Fig. 2B–D**). To confirm that the increase in RNA levels was due to active replication, infection was compared to treatment with rupintrivir. Viral RNA measured at 1 dpi was already greater than 2,000-fold for PV, or 20-fold for EV-D68 and EV-A71, above RNA levels measured in rupintrivir-treated samples (**Fig. 2B–D**). Of note, PV replicated more robustly than EV-D68 and EV-A71; PV RNA peaked at over a 100-fold increase from 1 dpi. Consistent with a higher replication capacity, PV produced a higher amount of viral progeny as quantified by plaque assay on cell culture supernatants containing virus released from infected hSpO (**Fig. 2E**). In addition, compared to PV, EV-D68 and EV-A71 produced more than a two log lower viral titer, and release of new viral progeny from the organoids above input virus was not observed until later time points of infection (8d and 2-4d, respectively) (**Fig 2E–G**).

**Figure 2:**
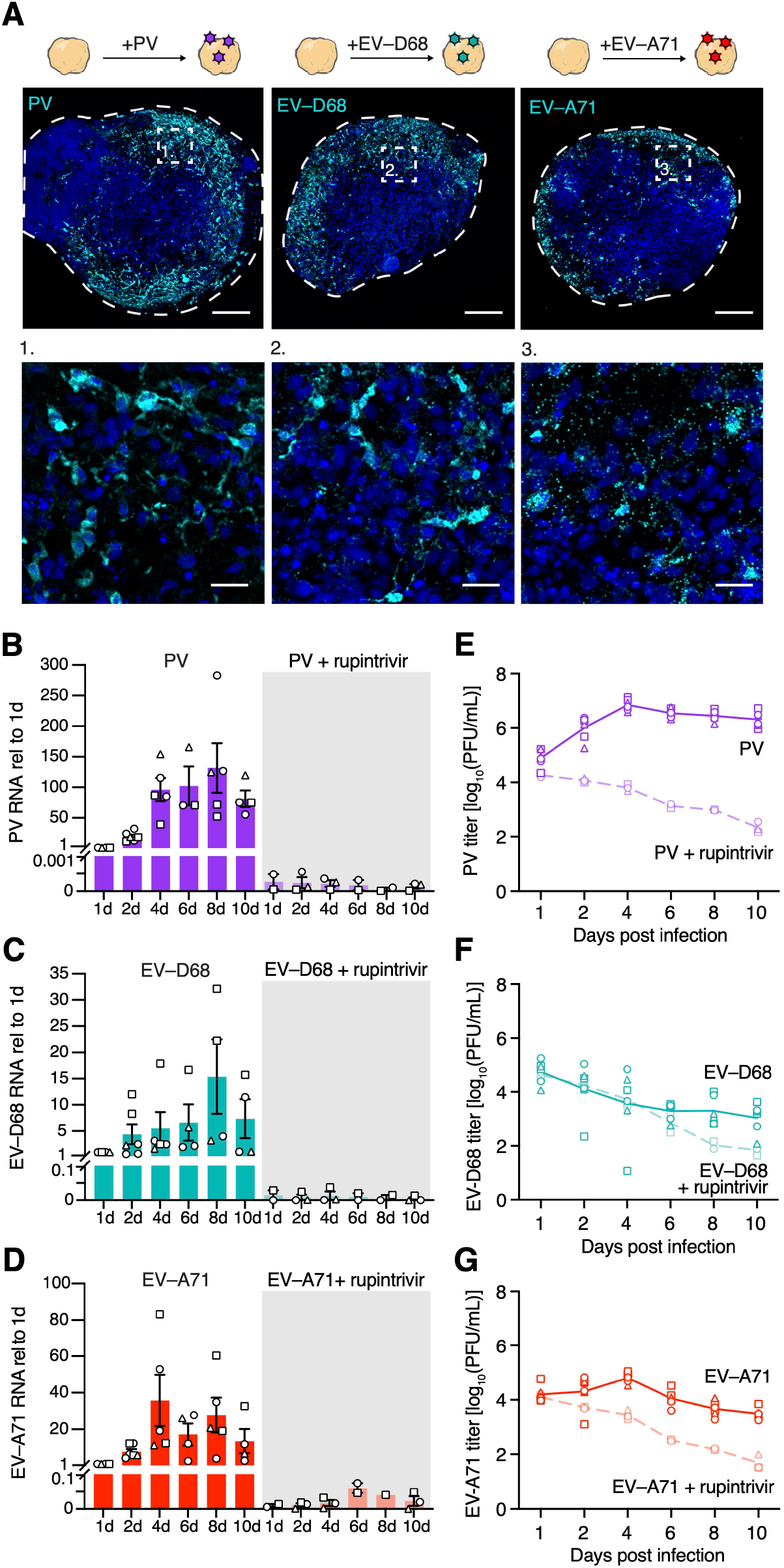
Productive infection of hSpO by polio and non-polio enteroviruses. **(A)** Schematic illustrating infection of hSpO with polio and non-polio EVs and representative immunofluorescent images of hSpO infected with PV, EV-D68 IL, or EV-A71 at 2 days post infection. Sections were stained with a panEV antibody to detect PV, an anti-D68 VP1 antibody to detect EV-D68, and an anti-dsRNA antibody to detect EV-A71. Nuclei are labeled with Hoechst (blue). Scale bars, 200 μm. Inset, scale bars 25 μm. **(B-D)** Intracellular viral loads in hSpO after exposure to PV, EV-D68 IL, or EV-A71 in the presence or absence of rupintrivir treatment were determined by qRT-PCR. Two hSpO were pooled per data point. n= 54 PV infected hSpO, n= 30 PV+rup infected hSpO, n= 54 EV-D68 infected hSpO, n= 30 EV-D68+rup infected hSpO, n= 56 EV-A71 infected hSpO, n= 28 EV-A71+rup infected hSpO from 3 hiPS cell lines from 1 differentiation. Values represent mean ± SEM. **(E-G)** Quantification of extracellular viral titers from culture supernatants of infected hSpO as determined by plaque forming units (PFUs)/mL on H1Hela cells (PV) or RD cells (EV-A71, EV-D68 IL). Each datapoint is the average of two technical replicates of cell culture supernatants pooled from 3-4 hSpO. n= 124 PV infected hSpO, n= 45 PV+rup infected hSpO, n= 124 EV-D68 infected hSpO, n= 45 EV-D68+rup infected hSpO, n= 125 EV-A71 infected hSpO, n= 45 EV-A71+rup infected hSpO from 3 hiPS cell lines from 2 differentiations (1 differentiation for rupintrivir treated samples). Line graphs represent average.

We proceeded to examine the cellular immune response of hSpO to EV infection by assessing the secretion of a panel of 80 human cytokines and chemokines via a multianalyte Luminex-based assay (**Supplemental Figure 1A**). Despite the lack of microglia or other immune cells in this system, the levels of several proinflammatory cytokines were increased upon infection by all three EVs, including cytokines previously associated with EV neuropathogenesis such as IL-6, IL-8 and CCL5 (RANTES) (Feng et al., 2020; Lin et al., 2003; Liu et al., 2018; Luo et al., 2019). Notably, while type I IFNs play a major role in antiviral defense, hSpO infection with PV, EV-D68, or EV-A71 did not lead to strong increases of any of the type I (IFN-α2), type II (IFN-γ), or type III interferons (IFN-λ2) assayed.

### Single cell transcriptomic profiling of enterovirus-infected hSpO and investigation of EV tropism

To identify the cell types infected by different EVs, we performed single cell RNA-sequencing (scRNA-seq) (10x Chromium) of infected hSpO at 2 dpi (hSpO generated from two hiPS cell lines from four differentiations; **Fig. 3A, Supplemental Figure 2A**). We integrated all samples and visualized the data using the uniform manifold approximation and projection (UMAP) dimensionality reduction technique (**Supplemental Figure 2B**). Following quality control and filtering, we obtained transcriptomes for 34,184 cells from uninfected samples, 27,278 cells from PV infected samples, 14,609 cells from EV-D68 infected samples and 29,596 cells from EV-A71 infected samples (**Supplemental Figure 2C**). We identified cells in our hSpO samples corresponding to the major spinal cord cell types by transferring cell type labels based on previously annotated hSpO clusters (Andersen et al., 2020). These corresponded to a diversity of neuronal cell types, including a group of motor neurons expressing *PHOX2B* and *ISL1* (MN), four ventral-like clusters that showed expression of *GATA3, SHOX2, LHX1, PAX2* and *EVX1* (V2b, V2a, V1, V0), a group of dorsal-like cells expressing *NHLH1, ST18, LHX2* and *LHX9* (clusters d4/6 and d1/d5), as well as glial-like cells including cycling progenitors, astroglia and oligodendrocytes, expressing *MKI67, SPARCL1* and *SOX10*, respectively (**Fig. 3B, Supplemental Figure 2D**). We also examined cluster neurotransmitter identity and found that it matched previously described annotations that included glutamatergic (*SLC17A6*^+^), cholinergic (*SLC5A7*^+^ or *SLC18A3*^+^), glycinergic (*SLC6A5*^+^), and GABAergic (*GAD1*^+^, *GAD2*^+^ or *SLC32A1*^+^) cells (**Supplemental Figure 2D**).

**Figure 3:**
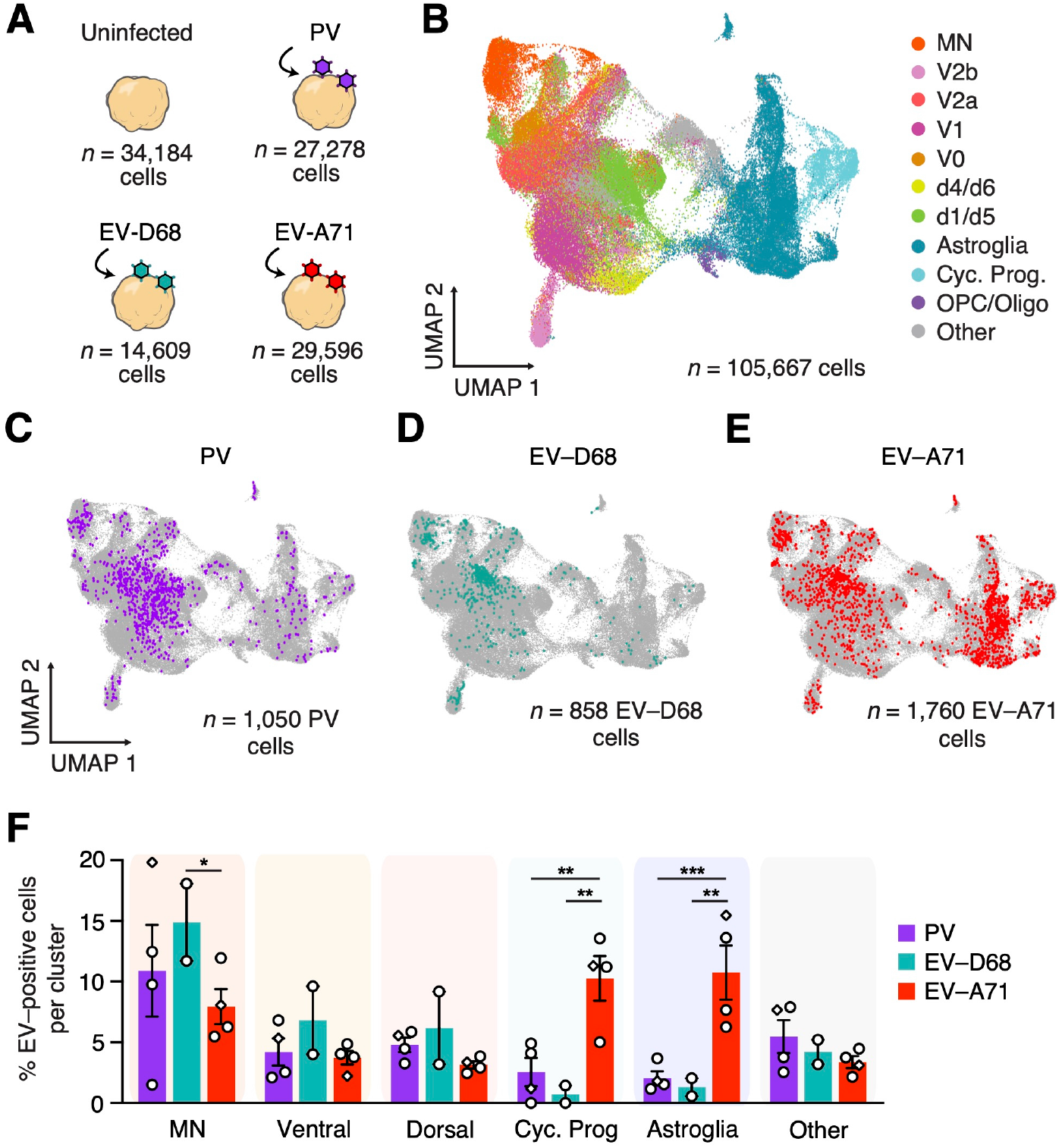
Single cell RNA sequencing of enterovirus infected hSpO to study cell tropism. **(A)** Schematic showing single cell RNAseq experimental design; hSpO were infected for 2 days prior to single cell dissociation. **(B)** UMAP visualization of single cell gene expression with cells colored by cell identity. All infected samples and uninfected controls are integrated and shown together. **(C-E)** UMAP plots highlighting positively infected cells per virus. **(F)** Percentage of infected cells per cell type in PV, EV-D68 and EV-A71 samples indicating enterovirus-specific tropisms. Each dot represents one single cell run, see Supplemental Figure 2. for information on all runs. Datasets represent mean ± SEM. P-values were determined by twoway ANOVA with Benjamini and Hochberg correction. *P<0.05, **P<0.01, ***P<0.001

Since EV transcripts are polyadenylated, we were able to map viral reads in the transcriptome to specific viruses. Viral RNA in hSpO cells could be due to active virus replication, or from abortive infections, ambient RNA from lysed cells, virus on the cell surface, or extracellular virus captured with single cells during GEM generation. To discriminate between cells supporting active virus replication versus those containing viral RNA due to other processes, we examined the distribution of viral reads across cells and observed clear bimodal distributions. In line with previously established methods (Kelly et al., 2022; Russell et al., 2018; Steuerman et al., 2018; Sun et al., 2020), we calculated kernel density estimates on these distributions and used the first local minima to generate cutoff thresholds to define two groups of cells: virus positive and virus negative (**Supplemental Figure 3A–C**). Abundant viral RNA was detected solely in the infected organoid samples and no viral transcripts were detected in uninfected hSpO.

We found that a subset of cells in each sample expressed high levels of viral RNA consistent with active viral replication (average of 3.85% for PV, 5.87% for EV-D68 and 5.95% for EV-A71 across all single cell sequencing runs) (**Supplemental Figure 3D**). When we compared the proportions of all cells in each cluster across infected and uninfected samples, we found that the overall diversity was similar and there were no obvious differences in the distribution of cell types (**Supplemental Figure 3E**). Next, to investigate what cell types were being preferentially infected, we compared the distribution of infected cells over the different cell clusters (**Fig. 3C–E**). We found that both PV and EV-D68 had a higher percentage of infection in MNs and other neuronal types (**Fig. 3F, Supplemental Figure 3F–G**). EV-A71 also infected cells in the MN cluster (**Fig. 3F, Supplemental Figure 3H**). However, in contrast to PV and EV-D68, EV-A71 exhibited specificity for infecting glial cell lineages (**Fig. 3F, Supplemental Figure 3H**). Consistent with a higher percentage of infected cells, the MN cluster contained a higher viral load for all EVs compared to the other neuronal cell clusters, and the cycling and astroglia clusters similarly had a higher average viral UMI for EV-A71 (**Supplemental Figure 3I**).

To further explore EV-specific tropism in hSpO and confirm the susceptibility of different cell types to EV infection, we performed live imaging of hSpO with labeled neurons or glia. To monitor EV infection by live imaging, we generated EVs encoding a fluorescent protein by introducing the coding sequence for enhanced green fluorescent protein (eGFP) or mNeon directly after the 5’UTR followed by a 2A cleavage site and the coding region of the viral polyprotein (**Supplemental Figure 4A**). To label neuronal or glial cells in hSpO we used either AAV-hSyn1-mScarlet to specifically label neurons or AdV-CMV-mCherry to label cells within the glial lineage (progenitors and astrocytes). Adenovirus (AdV) has been previously shown to preferentially infect glial lineage cells (Hansen et al., 2010; Qian et al., 2020), and we further confirmed this pattern of expression in hSpO by immunohistochemistry of AdV-CMV-mCherry-infected hSpO. Quantification of the co-localization of mCherry^+^ cells with the neuronal marker MAP2 or the glial marker SPARCL1, indicated that ∼80% of mCherry^+^ cells were of glial lineage, and fewer than 20% of them were neuronal (**Supplemental Figure 4B–C**).

Co-infection of fluorescent EVs with AAV-hSyn1-mScarlet or AdV-CMV-mCherry infected hSpO allowed us to monitor cell type-specific infection using live cell microscopy (**Fig. 4A**). We quantified the proportion of infected GFP/mNeon^+^ cells co-labeled with either AAV-hSyn1-mScarlet or AdV-CMV-mCherry. We found that ∼56% of all hSyn1-mScarlet labeled neurons were PV-mNeon^+^, ∼28% of all labeled neurons were EV-D68-GFP^+^ and ∼3% of all labeled neurons were EV-A71-mNeon^+^ (**Fig. 4B–C**). In line with our scRNA-seq results, we found that EV-A71 had a strong preference for infecting glial cells; ∼54% of all EV-A71-mNeon infected cells were AdV-CMV-mCherry^+^ while only ∼21% of all PV-mNeon infected cells or ∼3% of all EV-D68 infected cells were mCherry^+^ (**Fig. 4B–C**). Interestingly, upon infection of hSpO by EV-A71 we observed changes in the cellular morphology of astroglia, further highlighting the susceptibility of these cells to EV-A71 (**Fig. 4D**). In contrast, infection by PV or EV-D68 did not cause a change in the overall morphology of astroglia compared to uninfected hSpO (**Fig. 4D**). Importantly, we also observed a reduction in PHOX2B^+^ neuron numbers across 10 days of EV infection for all tested EVs (**Supplemental Figure 5A–B**), and this decrease in motor neurons was much more rapid upon PV infection. Taken together, these experiments confirm cell type specificity of EV infection in hSpO.

**Figure 4:**
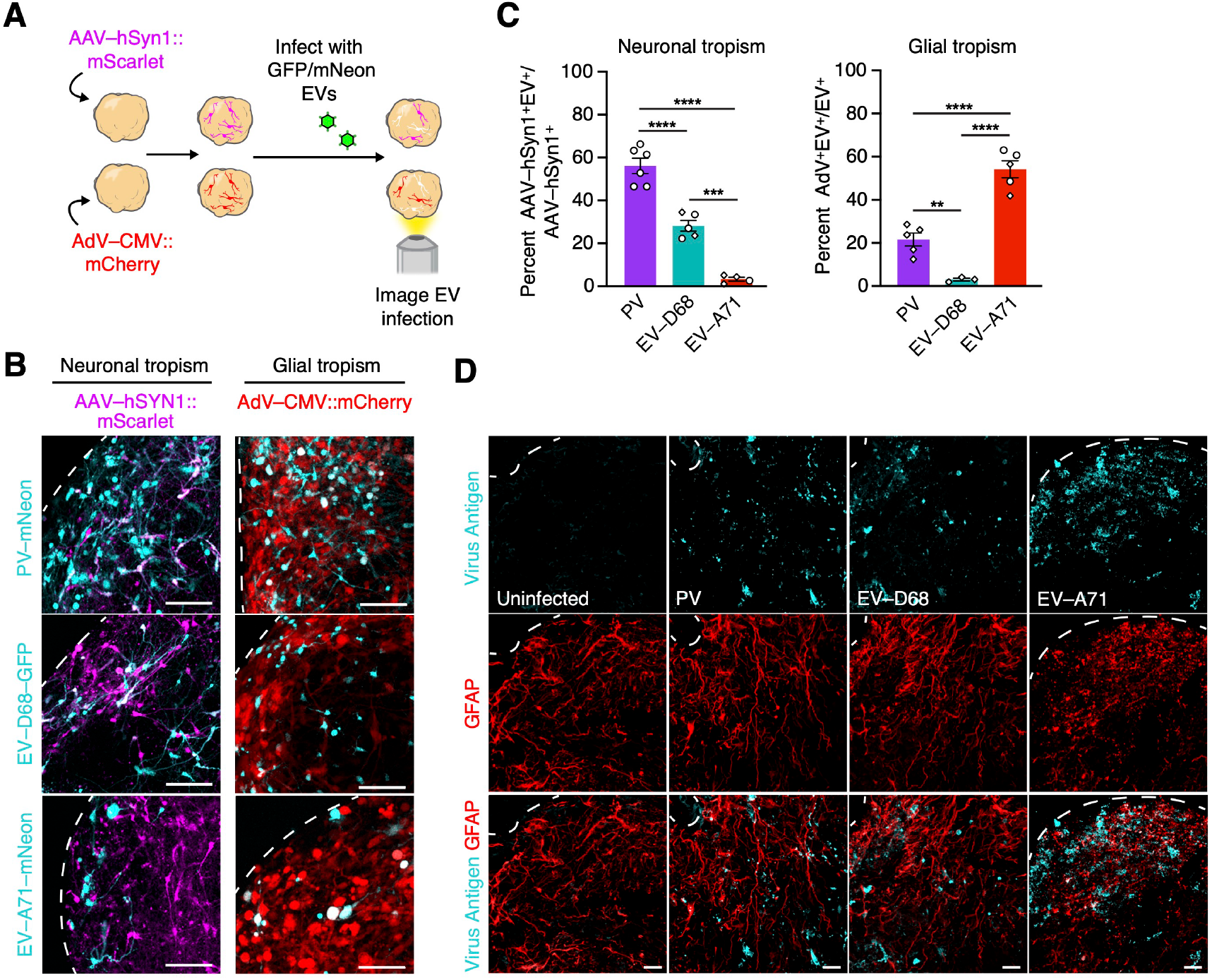
Glial and neuronal EV tropism. **(A)** Schematic showing experimental setup. hSpO were infected with either AAV-hSyn1-mScarlet (neuronal tropism) or AdV-CMV-mCherry (glial tropism) followed by infection with either PV-mNeon, EV-D68-GFP, or EV-A71-mNeon. **(B)** Representative images of live hSpO showing co-localization of EV-GFP/mNeon^+^ cells with either AAV-hSyn1-mScarlet or AdV-CMV-mCherry. Scale bar, 100 μm. **(C)** Quantification of EV-infected cells that colocalized with either AAV-hSyn1-mScarlet^+^ cells or AdV-mCherry^+^ cells showing neuronal or glial tropisms respectively. n= 11 PV infected hSpO from 2 hiPS cell lines from 1 differentiation, n= 8 EV-D68 infected hSpO from 2 hiPS cell lines from 2 differentiations, n= 9 EV-A71 infected hSpO from 2 hiPS cell lines from 2 differentiations. Values represent mean ± SEM. P-values were determined by one-way ANOVA adjusted with Benjamini-Hochberg, **P < 0.01, ***P < 0.001, ****P < 0.0001. **(D)** Representative GFAP immunostainings of hSpO infected with PV, EV-D68 or EV-A71 at 4 days post infection. Sections were stained with a panEV antibody to detect PV, anti-D68 VP1 antibody to detect EV-D68, and anti-dsRNA antibody to detect EV-A71. GFAP is labeled in red, virus is labeled in cyan. Scale bar, 25 μm.

### EV-mediated cell damage

The unique changes we observed in the glia and neurons upon EV infection prompted us to look more closely at the cellular features of infection. Intriguingly, live imaging of hSpO infected with fluorescent EVs revealed damage to infected cells and differences in the morphology of virus-induced damage over the course of 48 hours of infection. We categorized these damage phenotypes into two categories: when the cell body remained intact, and when the morphology of the cell body changed or was disrupted (**Fig. 5A**). Within infected cells where the cell body remained intact, we observed either no change in the shape of the soma (intact), or a potential decrease in the overall fluorescent intensity within the cell body (GFP decrease). For cells where the morphology of the cell body changed, we observed cells that either broke up into multiple smaller intact membrane-bound bodies (fragmented), or cells where the cell body would round up (rounded up). For each EV we quantified the proportion of virus infected cells that fell within these different damage bins. For PV infection, we found that greater than 50% of all infected cell bodies remained intact (**Fig. 5B, Supplemental Figure 5C**). Both EV-D68 and EV-A71 infection led to a markedly higher number of infected cells with disrupted cell bodies, with fragmented cell bodies representing greater than 60% of all infected cells for EV-D68 and rounded up cell bodies representing the main phenotype of cell damage for EV-A71 infected cells (**Fig. 5B, Supplemental Figure 5C**).

**Figure 5:**
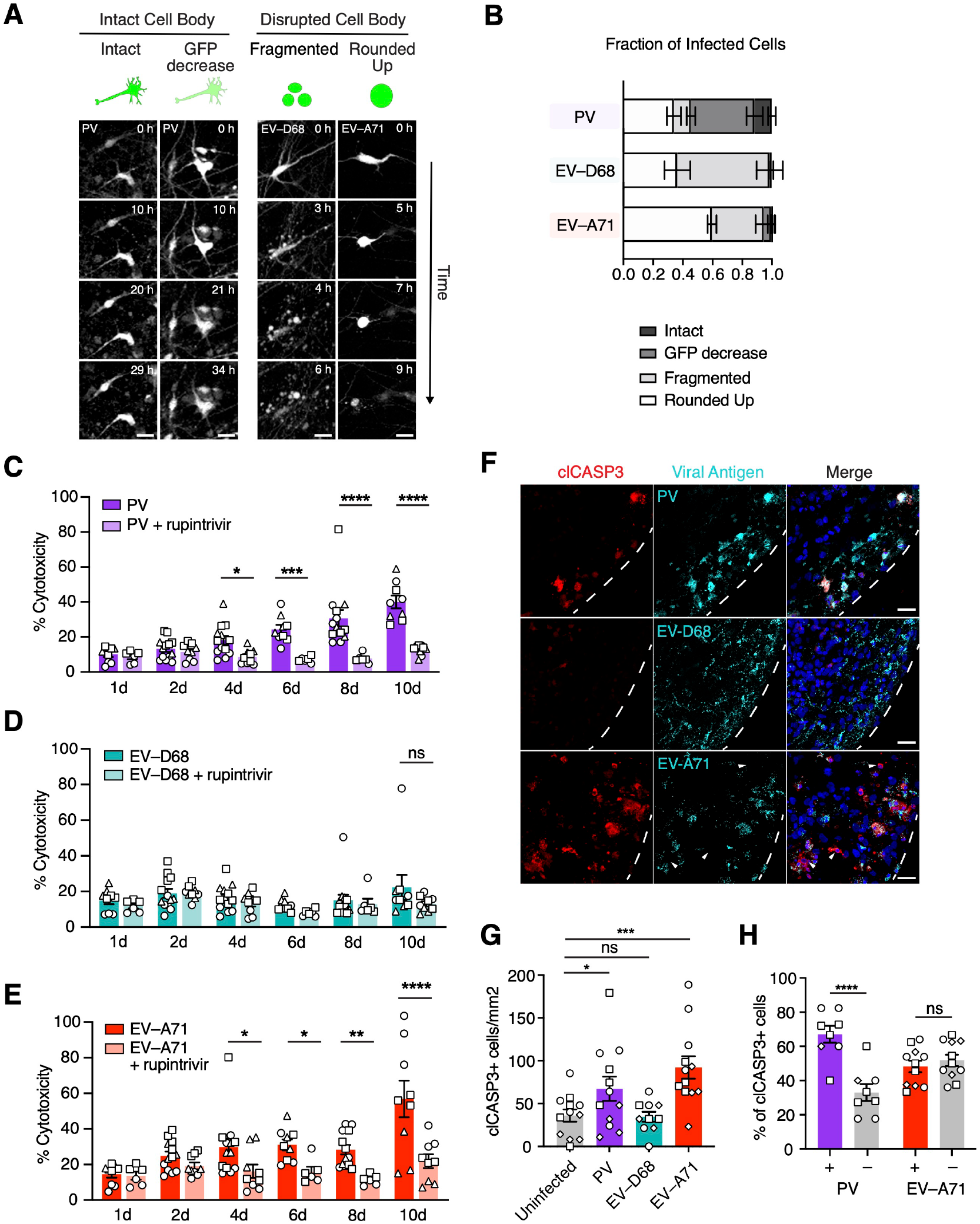
EV mechanisms of cell damage. **(A)** Representative live image sequences showing modes of cell damage after EV infection. Scale bars, 25 μm. **(B)** Stacked bar graphs showing the modes of damage per EV as percent. See Supplemental Figure 5C for individual data points. n= 9 PV infected hSpO from 2 hiPS cell lines from 1 differentiation, n= 7 EV-D68 infected hSpO from 2 hiPS cell lines from 2 differentiations, n= 5 EV-A71 infected hSpO from 2 hiPS cell lines from 2 differentiations. Values represent mean ± SEM. **(C-E)** Bar graphs showing percent cytotoxicity after infection of hSpO with C) PV, D) EV-D68, or E) EV-A71 as measured by amount of LDH released into cell culture supernatant at indicated time points in the presence or absence of 1 μM rupintrivir treatment. n= 66 PV infected hSpO, n= 45 PV+rup infected hSpO, n= 66 EV-D68 infected hSpO, n= 45 EV-D68+rup infected hSpO, n= 66 EV-A71 infected hSpO, n= 45 EV-A71+rup infected hSpO from 3 hiPS cell lines from one differentiation. Values represent mean ± SEM. P-values were determined by two-way ANOVA adjusted with Benjamini-Hochberg, *P < 0.05, **P < 0.01, ***P < 0.001, ****P < 0.0001. ns, non-significant. **(F)** Representative images of cleaved caspase 3 positive cells at 2 dpi of PV, EV-D68, or EV-A71 infected hSpO. Sections were stained with a panEV antibody to detect PV, or an anti-dsRNA antibody to detect EV-D68 and EV-A71. Nuclei are labeled with Hoechst (blue). White arrows denote uninfected cleaved caspase 3 positive cells. Scale bars, 25 μm. **(G)** Number of cleaved caspase 3 positive cells per hSpO area per virus. n= 12 uninfected hSpO, n= 12 PV infected hSpO, n= 10 EV-D68 infected hSpO, n= 12 EV-A71 infected hSpO from 3 hiPS cell lines from one differentiation. Values represent mean ± SEM. P-values were determined by ordinary one-way ANOVA adjusted with Benjamini-Hochberg, *P < 0.05, ***P < 0.001. **(H)** Fraction of cleaved caspase 3 positive cells that are EV positive or negative in PV or EV-A71 infected hSpO to examine cell autonomous versus non autonomous cell death. n= 12 PV infected hSpO, n= 12 EV-A71 infected hSpO from 3 hiPS cell lines from 1 differentiation. Values represent mean ± SEM. P-values were determined by two-way ANOVA adjusted with Benjamini-Hochberg, ****P < 0.0001.

To investigate damage beyond infected cells and assess overall damage to infected hSpO we measured the total amount of cell death over the course of 10 days of infection by measuring lactate dehydrogenase (LDH) release into the cell culture supernatant. LDH is a cytoplasmic enzyme that is released extracellularly upon loss of cellular membrane integrity. PV infection led to an increasing amount of damage over the course of infection with a peak cytotoxicity of ∼40% at 10 dpi (**Fig. 5C**). Upon EV-D68 infection we observed very minimal LDH release with only a 1.7-fold increase above rupintrivir treated spheres at 10 dpi (**Fig. 5D**). Surprisingly, we observed an even larger cytotoxic effect over the course of EV-A71 infection with a peak cytotoxicity of ∼60% at 10 dpi (**Fig. 5E**).

To further explore damage to hSpO, we assessed the presence of markers of apoptosis, a form of cell death previously shown to be induced upon EV infection (Lai et al., 2020; Rhoades et al., 2011). Apoptosis activation was monitored by immunostaining for cleaved caspase 3 (clCASP3) at 2 dpi (**Fig. 5F**). In line with the observed increase in cytotoxicity, both PV and EV-A71 infection significantly increased the number of clCASP3^+^ cells above uninfected hSpO (**Fig. 5G**). We infrequently observed clCASP3^+^ EV-D68 infected cells, and the total number of clCASP3^+^ cells was not significantly increased upon EV-D68 infection at 2 dpi (**Fig. 5F-G**). To investigate if cellular damage was occurring beyond infected cells, we next quantified the percentage of clCASP3^+^ cells that co-localized with viral antigens. Upon PV infection, the majority of apoptotic cells in infected hSpO co-localized with viral antigens (**Fig. 5H**). In contrast, EV-A71 infection led to an increase in clCASP3^+^ positive cells in both infected and uninfected cells (**Fig. 5H**), suggesting that EV-A71 infected cells can likely exert damage to non-infected cells in a non-cell autonomous manner.

## Discussion

Infection by AFM-causing EVs of the CNS has been primarily investigated through the use of animal models and cell systems. However, mice are refractory to infection by human EVs necessitating the use of transgenic mice models and other strategies including the use of mouse-adapted virus strains and infection of immunodeficient or neonatal mice, which limits their translational relevance (Racaniello, 2006; Vermillion et al., 2022; Wang and Yu, 2014). Therefore, there is a need for human cellular models of infection that recapitulate the cellular diversity, complex cell-cell interactions and circuit assembly of the spinal cord and associated periphery. In this study, we leveraged hiPS cell-derived human organoids and an assembloid model that recapitulates a functional human motor-neuron circuit (Andersen et al., 2020) to investigate the neuropathogenesis of polio and non-polio EVs. PV, EV-D68 and EV-A71 are three divergent neurotropic EVs that utilize different receptors and have differing primary sites of infection, yet they can all cause AFM. However, the mechanisms by which EV infection leads to paralysis are poorly understood. Here we found that infection of assembloids by all three EVs resulted in cessation of spontaneous muscle contraction that could be prevented by treatment with an antiviral drug. Our results recapitulate neuropathologic features observed in human infections supporting hSpO and assembloids as a relevant model system to study AFM.

We systematically compare infection by divergent neurovirulent EVs in our hSpO model. We show that PV, EV-D68 and EV-A71 productively infect and replicate within hSpO. However, we found that PV exhibited a higher replication capacity and reached a higher viral load compared to infection with EV-D68 and EV-A71. Single cell profiling of infected hSpO revealed tropism differences between the studied EVs. All viruses infected motor neurons, but while PV and EV-D68 mainly infected motor neurons and other neuronal types, EV-A71 preferentially infected astroglia. Using reporter viruses, we found evidence that the mechanisms by which these EVs cause damage may be distinct. EV-D68 and EV-A71 caused drastic morphological changes to individual infected cells causing them to fragment or round up, while a higher number of PV infected cells remained intact during long-term live-imaging. Despite this fragmentation for EV-D68 of initially infected cells, prolonged infection did not lead to measurable increases in cell death. In contrast, EV-A71 and PV infection resulted in strong cytotoxicity measured by LDH release and an apoptosis assay. Whereas apoptosis caused by PV infection was restricted to infected cells, EV-A71 caused apoptosis in both infected and surrounding uninfected cells. Astrocyte infection by EV-A71 could contribute to this indirect effect (Guttenplan et al., 2021). Altogether, our results suggest that divergent cell tropism and modes of cellular damage following EV infection converge on neuronal dysfunction leading to paralysis.

Our results are consistent with previously published studies using stem cell derived human spinal cord organoids and other multicellular systems to model EV infection. *In vivo* studies in mice and non-human primates showed a similar tropism for astrocytes by EV-A71 (Feng et al., 2016; Zhu et al., 2025). A recent study employing human spinal cord organoids similarly described motor neuron infection by EV-A71, but because infection was performed early after differentiation the organoids lacked astroglia (Chooi et al., 2024). Prior work with EV-D68 infection of spinal cord organoids also noted the apparent lack of appreciable cytopathic effect (Aguglia et al., 2023). However, in these previous studies, the functionality of the neurons was not tested. Importantly, our model recapitulates the cell diversity of the developing human spinal cord/hindbrain and includes functioning human neural circuits representing a significant advance beyond prior model systems.

While enteroviruses are the most well-known causes of AFM, AFM has been described for other neurovirulent viruses including West Nile virus (Fratkin et al., 2004; Nikolic et al., 2025; Sejvar et al., 2005). Beyond AFM, multiple viruses that infect and cause symptoms in the CNS enter the CNS via the spinal cord (Koyuncu et al., 2013) emphasizing the need for a physiological model of infection. We expect that our model can be broadly applied to study the neuro-pathogenesis of other viruses and can have utility for the development of antiviral therapies.

### Limitations of the study

This work also presents with limitations common to *in vitro* systems. Spinal cord organoids and assembloids represent a developmental model of neuromuscular connections; approaches aiming to incorporate more mature cell types such as myelinating oligodendrocytes could more faithfully assess cellular and functional responses. Similarly, neural organoids lack microglia and infiltrating immune cells. Studies aiming to incorporate these cell types through engraftment or transplantation would be beneficial in the future. Finally, for this study we focused on one strain per EV. While for PV all wild type strains are associated with paralysis, it is possible that for emerging non-polio enteroviruses such as EV-A71 and EV-D68 their ability to cause neuropathogenesis differs. A recent study observed differences in neurotropism and infectivity between two different EV-D68 strains (Aguglia et al., 2023; Dabilla et al., 2025). Future work could incorporate additional strains to better understand what underlies these differences.

## Acknowledgements

This work was supported by National Institutes of Health (NIH) grants R01-AI169467 (S.P.P. and J.E.C.), NIH T32 AI00732 (to C.E.P.), NIH T32G Stanford Cellular and Molecular Biology Training Program M007276 (L.V.), the National Science Foundation (GRFP Fellowship grant DGE-1656518 to C.E.P), the Idun Berry Postdoctoral Fellowship (J.A.), an Edward Mallinckrodt Jr Foundation award (J.A.), the National Institute of Neurological Disorders and Stroke F31 (5F31NS135955-02 to T.P.), the Stanford Brain Organogenesis Program in the Wu Tsai Neuroscience Institute and Bio-X (S.P.P.), Kwan Funds (S.P.P.), Senkut Funds (S.P.P.), and the Mann Foundation (S.P.P.). S.P.P. is a Chan Zuckerberg Initiative (CZI) Ben Barres Investigator and a CZ BioHub Investigator. J.E.C. is a Burroughs Wellcome Fund Investigator in the Pathogenesis of Infectious Disease. The funders had no role in the study design, data collection and analysis, decision to publish or preparation of the manuscript.

## Author contributions

J.A. and S.J.Y. performed hiPS cell differentiations and myoblast differentiations, and C.E.P assembled all hSpO-hSkM. C.E.P. and M.L. conducted assembloid and live imaging experiments and contributed to the analysis. C.E.P., L.V., and W.Q. generated and propagated all viruses used in these studies including fluorescent reporter viruses. C.E.P., J.A. and L.V. performed infection experiments, plaque assays and LDH assays. C.E.P., J.A., and M.V.T. performed immunostainings and C.E.P and J.A. performed confocal imaging. C.E.P., J.A., M.V.T., and X.C. performed the single cell transcriptomics experiments and C.E.P., J.A., and N.T contributed to the analysis. C.E.P and T.P. performed confocal data quantifications and analyses. C.E.P., J.A., M.L., J.E.C and S.P.P designed experiments and interpreted the experimental data. C.E.P., J.A., J.E.C and S.P.P. conceived the project and wrote the manuscript with input from all authors. S.P.P., J.E.C. and J.A. supervised the work.

## Materials and Methods

### Cells and Reagents

RD (CCL-136) cells and H1-Hela (CRL-1958) cells were obtained from the American Type Culture Collection (ATCC). Cell lines were cultured in Dulbecco’s modified Eagle’s medium (DMEM) with high glucose (Thermo Fisher Scientific, 11995-040) supplemented with 1× penicillin-streptomycin (Thermo Fisher Scientific, 15140163) and 10% heat-inactivated fetal bovine serum (Sigma Aldrich).

### Pluripotent stem cell culture and hSpO differentiation

The hiPS cell lines used in this study were validated using standard methods as previously described (Pasca et al., 2011; Sloan et al., 2018). A total of four hiPS cell lines derived from fibroblasts collected from four healthy subjects were used for experiments (see **Supplemental Table 1** for details of the hiPS cell lines used for each experiment). hiPS cell lines were maintained and passaged as previously described (Andersen et al., 2020; Yoon et al., 2019). Briefly, hiPS cells were maintained in human vitronectin-coated (VTN-N, Life Technologies, A14700) 6-well plates in Essential 8 medium (Life technologies, A1517001). Cells were incubated with 0.5 mM EDTA for 7 minutes at room temperature for passage, and resuspended and distributed in new 6-well plates in Essential 8 medium. Cultures were tested and maintained mycoplasma free. Approval for using these lines was obtained from the Stanford IRB panel and informed consent was obtained from all subjects.

The generation of hSpO from hiPS cells was performed using AggreWell 800 (STEMCELL Technologies, 34815) as previously described (Andersen et al., 2020) (Yoon et al., 2019). Following single cell dissociation using Accutase (Innovate Cell Technologies, AT-104) at 37 °C for 7 min, approximately 3 x 10^**6**^ single cells were added per AggreWell 800 well in Essential 8 medium supplemented with the ROCK inhibitor Y-27632 (10 µM, Selleckchem, S1049). Cells were then centrifuged at 100*g* for 3 min and incubated at 37 °C with 5% CO2. After 24 h, aggregated cells were collected from each microwell by firmly pipetting medium in the well and transferred into either ultra-low-attachment plastic dishes (Corning, 3262) or polyhema-treated dishes in Essential 6 medium supplemented with dual SMAD inhibitors dorsomorphin (2.5 µM, Sigma-Aldrich, P5499) and SB-431542 (10 µM, Tocris, 1614). For the first six days medium was replaced daily. From day 4 to day 18 medium was supplemented with the WNT activator CHIR 99021 (3 µM; Selleckchem, S1263). On day 6, hSpO were transferred to neural medium supplemented with RA (0.1 µM; Sigma-Aldrich, R2625), EGF (20 ng ml−1; R&D Systems) and FGF-2 (10 ng ml−1; R&D Systems), and addition of smoothened agonist (SAG, 0.1 µM; Millipore, 566660) from day 11. From day 7, the medium was then changed every other day. On day 19, hSpO were transferred to neural medium with N-2 supplement (Life Technologies, 17502048), BDNF (20 ng ml−1, Peprotech), IGF-1 (10 ng ml−1; Peprotech, 100-11), L-Ascorbic Acid (AA, 200 nM; Wako, 321-44823) and cAMP (50 µM; Sigma-Aldrich, D0627). On days 19, 21 and 23 the Notch pathway inhibitor DAPT (2.5 µM; STEMCELL technologies, 72082) was added. From day 43 onward, the medium was changed every four to five days.

### Culture of hSkM

Human skeletal myoblasts (hSkM) were obtained from Thermo Fisher Scientific (A11440, Lot# 2540962) and maintained in an undifferentiated state with Skeletal Muscle Cell Growth Medium (ready to use; Promocell, C-23060) in 10-cm plates (Primaria Cell Culture Dish, Corning). Medium was changed every 2–3 days. hSkM at ∼80% confluency were passaged using Trypsin (Trypsin-EDTA, 0.25%, phenol red; Life Technologies), and passages 1 to 4 were used for experiments.

### Generation of 3D hSkM

3D hSkM cultures were generated as previously described (Andersen et al., 2020). Here, 30,000 hSkM were resuspended in 10 µL of Geltrex™(Life Technologies) and placed in small silicone wells (Ibidi, 80409) inside 6-well tissue culture plates (Corning). After a 30-minute incubation, 4 mL of Skeletal Muscle Cell Growth Medium was added to the wells. Silicone wells containing hSkM were placed into 6-well ultra-low attachment plates on the next day, and media changes were performed every 2–3 days after that. Medium was changed to Skeletal Muscle Cell Differentiation Medium after 14 days, to allow for differentiation of hSkM with medium changes every 2–3 days.

### Generation of hSpO-hSkM

To generate assembloids, 3D hSkM that had been in differentiation medium for at least 14 days were placed on top of 6-well cell culture inserts (0.4 µm pore size; Corning, 353090) in wells containing 2 mL of DMEM/F12 medium supplemented with 1% Non-Essential Amino Acids (NEAA; Life Technologies), 1% Insulin-Transferrin-Selenium (ITS; Life Technologies), 1% penicillin and streptomycin (Life Technologies), L-Ascorbic Acid (AA, 200 nM; Wako) and cAMP (50 µM; Sigma-Aldrich). hSpO organoids at days 28-32 of differentiation were then placed on the inserts in contact with 3D hSkM. hSpO-hSkM assembloids were allowed to interact for a minimum of 10 days before being used in experiments. Between 1-3 hSpO-hSkM assembloids were maintained per insert, and half medium changes were performed every other day. Assembloids were screened prior to the initiation of experiments, and only assembloids that displayed spontaneous contractions at baseline were included for EV treatment; this corresponded to the inclusion of 56 out of 121 assembloids screened.

### Viruses

EV-D68 2014 outbreak isolate (US/IL/14-18952; NR-49131) was provided by BEI Resources (National Institute of Allergy and Infectious Diseases (NIAID), National Institutes of Health (NIH)). EV-D68 was propagated and titred by plaque assay on RD cells at 34°C. The PV-1 Mahoney strain was a gift from H. Ploegh and was propagated and titred on H1-HeLa cells. EV-A71 (BrCr strain) was obtained from ATCC and was propagated and titred by plaque assay on RD cells.

### Construction of reporter viruses and infectious clones

Fluorescent reporter EVs were generated by introducing the coding sequence for enhanced green fluorescent protein (eGFP) or mNeon directly after the 5’UTR followed by a 2A cleavage site and the coding region of the viral polyprotein. This design was based on a published strategy to generate an infectious clone for the related enterovirus CV-B3 (Lanke et al., 2009).

The plasmid pPVM-2A144-DsRed, which encodes the PV genome with the insertion of the DsRed coding sequence after amino acid 144 of 2A with an additional 9 nt coding for three spacer amino acids on each side (Teterina et al., 2010), was used as a template for the construction of plasmid pPVM-mNeon-GPI. The reporter virus construct was assembled from three fragments. The pPVM-2A-144-DsRed clone was PCR-amplified into two fragments by Phusion High-Fidelity DNA Polymerase using the primer pairs: F1-F: 5’-CGAGGGTGGTGGTGGAAGTTGCCTG-3’ and F1-R: 5’-CCATGGCTTCTTCTTCGTAGGCATACAAGTCTCTAATGTCTG-3’; F2-F: 5’-CTACGAAGAAGAAGCCATGGAACAAGGCCTCACCAATTAC-3’ and F2-R: 5’-TGAGCGCAACGCATCGAAGATTCCGAG-3’. The third fragment containing the reporter gene insertion, an mNeon reporter gene followed by a farnesyl tag and the PV 2C cleavage site (MEALFG) between PV 5’ UTR and the polyprotein, and the rest of the virus genome was synthesized with IDT. Three fragments contained 20 bp overlaps at both ends and were assembled into the final construct PV-mNeon-GPI using Gibson Assembly (New England Biolabs).

The EV-A71 4643 infectious clone, which encodes cDNA for the EV-A71 Taiwan/4643/98 strain under the control of a T7 promoter was produced by the lab of Jen-Ren Wang (Huang et al., 2012). The full-length EV-A71 4643 infectious clone was used as a template to insert an mNeon reporter gene between the EV-A71 5′ UTR and the polyprotein. Following the reporter gene, the amino acids AITTL were added by replacing the original polyprotein start codon to introduce an EV-A71 2A cleavage site (AITTLG). To generate the reporter virus construct, the EV-A71(4643) infectious clone was linearized using BlpI and AflII restriction enzymes, and a 1344 bp fragment containing the reporter gene insertion and the rest of the virus genome was synthesized with IDT. Subsequently, the two fragments were assembled using Gibson Assembly (New England Biolabs).

EV-D68 IL infectious clone pUC-49131, which contains cDNA from EV-D68 US/IL/14-18952 under the control of a T7 promoter was provided by BEI Resources (National Institute of Allergy and Infectious Diseases (NIAID), National Institutes of Health (NIH), NR-52011). The full-length EV-D68 pUC-49131 infectious clone was used as a template to insert a GFP reporter gene between the EV-D68 5′ UTR and the polyprotein. Following the reporter gene, the amino acids AAAEALFQ were added by replacing the original polyprotein start codon to introduce a 2A cleavage site. To generate the reporter virus construct, the EV-D68 pUC-49131 infectious clone was linearized using the MluI-HF restriction enzyme, and a 2033 bp fragment containing the reporter gene insertion and the rest of the virus genome was amplified by PCR. Subsequently, the two fragments were assembled using Gibson Assembly (New England Biolabs).

Engineered infectious cDNAs were then linearized by restriction digestion using the restriction enzyme MluI-HF (New England Biolabs) for the EV-A71 infectious clone, the restriction enzyme SalI-HF (New England Biolabs) for the EV-D68 infectious clone, and the restriction enzyme EcorI-HF (New England Biolabs) for the PV clone. Linearized cDNA was purified using QIAquick PCR purification kit (Qiagen), and in vitro transcribed using the MEGAscriptTM T7 Transcription Kit (Invitrogen). RNA was purified using the RNeasy MinElute Cleanup Kit (Qiagen), and RNA quantity and quality were assessed using a Bioanalyzer 2100 RNA Nano Kit (Agilent).

To generate PV-mNeon and EV-D68-GFP infectious virus, H1-HeLa^**+CDHR3**^ (WT) cells in electroporation buffer (Teknova) were electroporated with the indicated purified viral RNAs using the square wave protocol (120 V; 1.5 ms pulse; 10 pulse; 1.5 s interval; 1 mm) on a Gene Pulser XcellTM electroporation system (Bio-Rad). Cells and supernatants were harvested 1 day after infection for PV-mNeon and 10 days after infection for EV-D68-GFP. The titer of the PV-mNeon viral stock was determined by plaque assay on H1-Hela cells. To determine the titer of the EV-D68-GFP viral stock, RD cells seeded in 6 well plates were incubated with EV-D68-GFP for 2 h at 34°C. After virus adsorption, viral inoculum was removed and replaced with 3mL of overlay medium and further incubated at 34°C. Overlay medium consisted of 2X DMEM supplemented with 2× penicillin-streptomycin (Sigma), and 20% heat-inactivated fetal bovine serum (Sigma), combined with 3% low melting temperature agarose. At 4 dpi fluorescent foci of infection were counted on an Incucyte S3 live-cell analysis system (Sartorius) and the viral titer was calculated as fluorescent focus units (FFU) per ml. To generate infectious EV-A71-mNeon virus, RD cells in electroporation buffer (Teknova) were electroporated with the purified viral RNA using the square wave protocol (120 V; 1.5 ms pulse; 10 pulse; 1.5 s interval; 1 mm) on a Gene Pulser XcellTM electroporation system (Bio-Rad). Cells and supernatants were harvested after 4 days. The titer of the viral stock was determined by plaque assay on RD cells.

### Enterovirus infection of hSpO-hSkM and hSpO

hSpO-hSkM were infected with 10^**4**^ plaque-forming units (PFU) of PV, 10^**5**^ PFU of EV-D68 or 10^**4**^ PFU of EV-A71 per assembloid at 37°C. Virus was added on top of the hSpO region on the cell culture insert in a volume of less than 3uL. A half medium change was performed at the time of infection and then performed every other day. For rupintrivir treated samples, 1µM rupintrivir was added to the assembloid at the time of infection and was re-added to the media every 2 days at the time of media change. For treatment, approximately 5uL of media containing rupintrivir was added on top of the cell culture insert to surround the assembloid.

hSpO were infected with 10^**4**^ PFU of PV, 10^**5**^ PFU of EV-D68 or 10^**4**^ PFU of EV-A71 per organoid at 37°C. For all experiments, hSpO were infected between days 76 and 125 of differentiation; ages of hSpO used for single cell experiments are listed below under Single cell dissociation. Each hSpO was placed in an individual well of a 24-well plate (low attachment or polyhema-treated) in 500 µL of fresh hSpO culture medium. hSpO were maintained in hSpO culture medium with virus until the indicated time point of collection. For timepoints greater than 5 days of incubation, an additional 500 µL media was added at 5 days post infection (dpi). To test for active RNA replication, 1µM rupintrivir was added to the spheres at the time of infection. Rupintrivir was re-added to the media every 2 days and at 5 dpi when additional media was added to the samples.

### Imaging of spontaneous contractions in hSpO-hSkM

Spontaneous muscle contractions were imaged under environmentally controlled conditions (37°C, 5% CO_**2**_) using a 10x objective in a Leica SP8 or Leica Stellaris 5 confocal microscope. Assembloids, still in transwells, were incubated in the environmentally controlled chamber for 20–30 minutes before imaging, and they were imaged for 2 minutes at a frame rate of 14.7 frames/sec. 1–3 fields were imaged per assembloid and each field was analyzed separately. Imaging was performed every 2 days, with the same fields imaged at each timepoint.

### Muscle contraction analysis

Muscle contraction of 3D hSkM was quantified by analyzing changes in pixel intensity over time using the automated, open-source ImageJ plugin MUSCLEMOTION (Sala et al., 2018) as previously described (Andersen et al., 2020) (https://github.com/l-sala/MUSCLEMOTION). MUSCLEMOTION speed of contraction was used for event detection. Event detection was performed using custom MATLAB routines, with events over 4 standard deviations counted as a contraction event. Each imaging field (931 µm by 931 µm in size) was divided into 4 subfields and the analysis was performed in each of the subfields individually. The median number of contractions from the four subfields of a field was then graphed.

### Cryopreservation and immunohistochemistry

Cryopreservation and immunocytochemistry of hSpO and hSpO-hSkM was performed as previously described (Andersen et al., 2020; Sloan et al., 2018). Briefly, organoids or assembloids were fixed in 4% paraformaldehyde (PFA in PBS, Electron Microscopy Sciences) for 2 hours. Following fixation, samples were washed twice with PBS, and cryopreserved by first placing in sucrose (30% sucrose in PBS for 24-48 hours or until samples sank to the bottom), and then embedding in 1:2, 30% sucrose: OCT (Tissue-Tek OCT Compound 4583, Sakura Finetek) and freezing. For immunocytochemistry, 16 µm thick sections were obtained using a cryostat (Leica). Cryosections were washed with PBS to remove excess OCT, blocked for 1 h at room temperature (10% normal donkey serum (NDS), 0.3% Triton X-100 diluted in PBS or 5% BSA, 0.02% sodium azide and 0.3% Triton-X-100 diluted in PBS), and incubated overnight at 4°C with primary antibodies in blocking solution. The primary antibodies used to detect the presence of the indicated proteins in this study are as described: dsRNA clone rJ2 (MABE1134; 1:50; Millipore), EV-D68 VP1 (GTX132313; 1:200; Genetex), enterovirus pan monoclonal antibody L66J (MA5-18206; 1:50; Thermo Fisher Scientific), Desmin (50-173-1004; 1:200; Thermo Fisher Scientific), cleaved caspase-3 (9661; 1:400; Cell Signaling Technology), GFAP (13-0300; 1,000; Thermo Fisher Scientific), PHOX2B (AF4940-SP; 1:2,000; R&D Systems), MAP2 (188 004; 1:5,000; Synaptic Systems) and SPARCL1 (AF2728; 1:300; R&D Systems). Next day, cryosections were washed with PBS and incubated with secondary antibodies for 1 h at room temperature. Alexa Fluor secondary antibodies (Life Technologies) were used diluted in blocking solution at 1:1,000. Following washes with PBS, nuclei were visualized with Hoechst 33258 (Life Technologies). Finally, slides were mounted for microscopy with cover glasses (Fisher Scientific) using Aquamount (Polysciences) and imaged on a Zeiss LSM 700 confocal microscope. Images were processed in ImageJ (Fiji) and are presented as projections from a z-stack.

### Real-time quantitative PCR - qPCR

For qPCR analysis of hSpO, 2 organoids were pooled per sample. mRNA was isolated using either the RNeasy Micro kit and RNase-Free DNase set (Qiagen) or the miRNeasy Micro kit (Qiagen). Template cDNA was prepared by reverse transcription using the SuperScript III First-Strand Synthesis SuperMix for qRT–PCR (Life Technologies). qPCR was performed using SYBR Green (Roche) on a CFX Connect Real-Time PCR Detection System (Biorad). Viral RNA levels were normalized to actin RNA levels.

The following primers (5′–3′ orientation) were used: actin-Fwd:

AGGTCTTTGCGGATGTCCACGT;

actin-Rev: CACCATTGGCAATGAGCGGTTC;

PV-Fwd: CAACCTCCCACTGGTGACTT;

PV-Rev: ATTTCCCCTGCTCAACCTTT;

EV-A71-Fwd; CCCTGAATGCGGCTAATCC;

EV-A71-Rev: ATTGTCACCATAAGCAGCCA;

EV-D68-Fwd: GGTGTGAAGAGTCTATTGAGC;

EV-D68-Rev: CACCCAAAGTAGTCGG

### Extracellular viral titer quantification by plaque assay

H1HeLa cells (for PV) or RD cells (for EV-D68 and EV-A71) were seeded in 6 well plates and incubated with the cell culture medium collected from PV, EV-D68 or EV-A71 infected hSpO for 2 h at 37°C (PV, EV-A71) or 34°C (EV-D68). After virus adsorption, inoculum was removed and replaced with 3mL of overlay medium and further incubated at the corresponding temperatures. Overlay medium consisted of 2X DMEM supplemented with 2× penicillin-streptomycin, and 20% heat-inactivated fetal bovine serum, combined with 3% NuSieve low-melting-temperature agarose (for EV-A71) or 2.4% Avicel (PV, EV-D68). Cells were fixed at 2 dpi (PV), 4-5 dpi (EV-D68), or 5-7 dpi (EV-A71), and stained with crystal violet. Cell culture medium from 3-4 organoids were pooled for each timepoint, the data represents the average of a technical duplicate.

### Single cell dissociation

hSpO were infected at day 62 (R1), day 80 (R2), day 98 (R3), or day 68 (R4) of differentiation (4 independent differentiations). After two days of infection, organoids were dissociated into single cells and processed on the same day. Five organoids for each condition were combined per dissociation. Dissociation of hSpO into single cells was performed as previously described (Birey et al., 2017; Pasca et al., 2015; Sloan et al., 2017; Sloan et al., 2018). Briefly, 5 hSpO per sample were chopped and incubated in 40 U/ml papain enzyme solution at 37°C for 90 minutes. After digestion, samples were washed with a protease inhibitor solution and gently triturated to achieve a single cell suspension and strained using a 70 µm Flowmi strainer (Bel-Art).

### Single cell gene expression - 10x Genomics

Single cell dissociated cells were resuspended in ice-cold PBS containing 0.02% BSA and loaded onto a Chromium Single cell 3′ chip (with an estimated recovery of 10,000 cells per channel). scRNA-seq libraries were prepared with the Chromium Single cell 3′ GEM, Library & Gel Bead Kit v3 (10x Genomics, PN: 1000075). Libraries were pooled and sequenced by Admera Health on a NovaSeq S4 (Illumina) or NovaSeq X Plus using 150 × 2 chemistry.

Demultiplexing, alignment, barcode and UMI counting and aggregation were performed using Cell Ranger (v6.0 with default settings). The human genome, Ensembl GRCh38.98.gtf, the PV1 Mahoney genome, NCBI Genome database accession NC_002058.3, EV-A71 BrCr genome, accession U22521.1, and EV-D68 genome, accession KM851230.1, were combined and used for alignment. Further analysis was performed using the R package Seurat (v5) from the Satija Lab (Butler et al., 2018). We excluded genes that were not expressed in at least three cells, and we excluded cells with less than 2,000 detected genes, as well as those with a mitochondrial content higher than 10%.

We normalized and scaled gene expression for each sample using a regularized negative binomial regression method (SCTransform function v1) (Hafemeister and Satija, 2019). We then integrated cells across samples to analyze the response to infection. To do so, we selected the 3,000 most variable genes across the datasets (SelectIntegrationFeatures function with nfeatures = 3,000) and performed a dimensionality reduction (RunPCA function). Harmony was run for batch correction (RunHarmony function) using both hiPS line and individual single cell sequencing run variables as arguments of “group.by.vars” (Korsunsky et al., 2019). UMAP embeddings (RunUMAP) were then obtained using the first 20 Harmony dimensions, and clusters were obtained by identifying shared nearest neighbors (FindNeighbors with dims 1:20) and running the FindClusters function with a resolution of 1.5.

### Cell Type Annotation

To annotate the identity of the cells in our samples we performed a label transfer in Seurat leveraging our previously published hSpO dataset as a reference (Andersen et al., 2020). Pairs of anchors between the two datasets were identified using the FindTransferAnchors function (dims 1:20) and the TransferData function was used to score each of the cells in the query dataset for similarity with labeled cell types in the reference dataset. Because the two datasets differed in age, with the newly generated dataset having used older organoids, we then further annotated cells initially labeled as progenitor or astroglia by label transfer into OPC/Oligo, cycling or astroglia lineages based on known cell-type specific markers. The OPC/Oligo cluster consisted of a small number of cells (less than 50 cells per sample for most runs) and were therefore not included in the subsequent analysis.

### Viral gene expression

To determine cells that were either positive or negative for each enterovirus, we determined a threshold by examining the distributions of viral genome expression across cells for each library, which followed a clear bimodal distribution (Kelly et al., 2022; Russell et al., 2018; Steuerman et al., 2018; (Sun et al., 2020). We calculated kernel density estimates on these distributions and used the first local minima to generate cutoff thresholds. Cells that fell within the lower peak were assumed to not be truly infected and were classified as virus negative cells. Cells that fell within the higher peak were categorized as virus positive.

### Live imaging

Prior to enterovirus infection, organoids were infected with either AAV1-hSYN1-mScarlet (Addgene, 131001; 9.5×10^**12**^ GC/ml) to label neurons or AdV-CMV-mCherry (adenovirus type 5; Vector Biolabs, 1767; 1×10^**10**^ PFU/ml) to label glial cells. For AAV infection, approximately 0.2 µl AAV was used for each organoid. Approximately 5 organoids were pooled in an eppendorf tube with minimal medium for about ∼2h, and 500 µl medium overnight. Organoids were transferred to 24 well plates the next day. For AdV infection, 5-10 organoids were pooled in a 24 well and infected with 1-2 µl of adenovirus in 1 mL of culture medium. Organoids were used in experiments after 3-4 weeks of AAV infection or 2–4 days of AdV infection.

For all live imaging experiments, AAV1-hSYN1-mScarlet or AdV-CMV-mCherry labeled organoids were infected in a synchronized manner with 10^**4**^ PFU PV-mNeon, 10^**5**^ PFU EV-D68-GFP or 10^**4**^ PFU EV-A71-mNeon per organoid. In this case, organoids were incubated with the corresponding PFU in 25 µL of fresh hSpO culture medium in an eppendorf tube for 2 hours at 37°C to allow for infection, after which the virus inoculum was removed, organoids were washed and plated in 96 well imaging plates (96-well µ-plate, Ibidi, 89621). One organoid was plated per well with 300 µL of fresh hSpO culture medium. Imaging started 1–3 hours post infection. Imaging was performed at 37°C with a 10× objective on a Leica Stellaris 5 confocal microscope with 5% CO_**2**_. hSpO organoids were imaged at an interval of 1h, and imaging lasted for 16-48h. For AdV imaging, due to strong mCherry fluorescence, a stringent cutoff was set to prevent bleeding into the 488 channel. Images are presented as projections from a z-stack with a z-interval of 5–6 µm. Imaging was corrected for drifting with StackReg plugin in Fiji (Thevenaz et al., 1998).

### Live imaging Quantification

For quantification of EV-infected cells that colocalized with AAV-hSyn1mScarlet, since AAV-hSyn1-mScarlet did not efficiently label all neuronal cells within the organoid, we manually counted the total number of mScarlet^**+**^ labeled neurons and then determined the number that were GFP/mNeon^**+**^. The total number of AAV-hSyn1-mScarlet^**+**^ cells was determined at the beginning timepoint (t0) of the imaging experiments. For quantification of the number of infected GFP/mNeon^**+**^ cells co-labeled with AdV-CMV-mCherry, the total number of GFP/mNeon^**+**^ cells was manually counted at a single timepoint during the imaging experiment (at approximately a quarter to halfway through an imaging session).

For quantification of the damage bins, EV-GFP/mNeon infected cells were identified within co-labeled hSpO at an early timepoint of the live imaging session (at approximately the 1/4^**th**^ point) and were surveyed for cells with cell bodies that could be tracked throughout the duration of the imaging session. On average, 15 EV-GFP/mNeon^**+**^ cells per area were inspected for damage and categorized into one of the four corresponding categories using ImageJ.

### Luminex-EMD Millipore Human 80 Plex

For Luminex analysis of hSpO, cell culture supernatants from 2 hSpO were pooled per sample. Supernatants from uninfected and infected hSpO were UV-inactivated at 300 mJ and stored at −80°C prior to analysis by Luminex. This assay was performed by the Human Immune Monitoring Center at Stanford University-Immunoassay Team. Kits were purchased from EMD Millipore Corporation, Burlington, MA., and run according to the manufacturer’s recommendations with modifications described as follows: H80 kits include 3 panels: Panel 1 is Milliplex HCYTA-60K-PX48. Panel 2 is Milliplex HCP2MAG-62K-PX23. Panel 3 includes the Milliplex HSP1MAG-63K-06 and HADCYMAG-61K-03 (Resistin, Leptin and HGF) to generate a 9 plex. The assay setup followed recommended protocol. Briefly, 25µl of the undiluted sample was mixed with antibody-linked magnetic beads in a 96-well plate and incubated overnight at 4°C with shaking. Cold and room temperature incubation steps were performed on an orbital shaker at 500-600 rpm. Plates were washed twice with wash buffer in a BioTek ELx405 washer (BioTek Instruments, Winooski, VT). Following one-hour incubation at room temperature with biotinylated detection antibody, streptavidin-PE was added for 30 minutes with shaking. Plates were washed as described above and PBS added to wells for reading in the Luminex FlexMap3D Instrument with a lower bound of 50 beads per sample per cytokine. Each sample was measured in duplicate. Custom Assay Chex control beads were purchased and added to all wells (Radix BioSolutions, Georgetown, Texas). Wells with a bead count <50 were flagged.

### LDH assay

Cell death was assessed by measuring levels of lactate dehydrogenase (LDH) released into the cell culture supernatant using LDH-Glo Cytotoxicity Assay (J2381; Promega). Luciferase readings were taken one hour after reagent addition and read using a GloMax 20/20 Luminometer. For each individual hiPS line and corresponding differentiation, a maximum LDH release control was determined by incubating uninfected hSpO with 2 µL of 10% Triton X-100 per 100µl media for 1 hour prior to collecting the cell culture supernatant. The average luminescence value of the Triton-X-100 treated samples was set as 100% for each line and data is graphed as percent cytotoxicity relative to this control.

### Immunohistochemistry Quantification

For quantification of the number of cleaved caspase 3 positive cells per area, images were acquired on a Zeiss LSM 700 confocal microscope, and fields of 125 x 125 µm were obtained for analysis. All fields were taken within 200 µm of the organoid edge in order to exclude the necrotic core. For each experimental condition, at least 2-6 fields per section and 2-3 16 µm thick sections per organoid were quantified (Pedrosa et al., 2020); dataset represents the average for each organoid. Images were quantified with Image J software.

For quantification of the number of PHOX2B positive cells per area, the total number of cells expressing PHOX2B per hSpO area (mm^**2**^) were counted in 16 µm thick cryosections of PV, EV-D68, or EV-A71 infected hSpO. PHOX2B signal was only counted if it overlapped with a nuclei (Hoechst) corresponding to a cell body in the section. 1-3 sections per organoid were quantified and averaged per organoid.

### Statistical analysis

GraphPad Prism 10 (GraphPad Software) was used to perform statistical analyses.

**Supplemental Figure 1:**
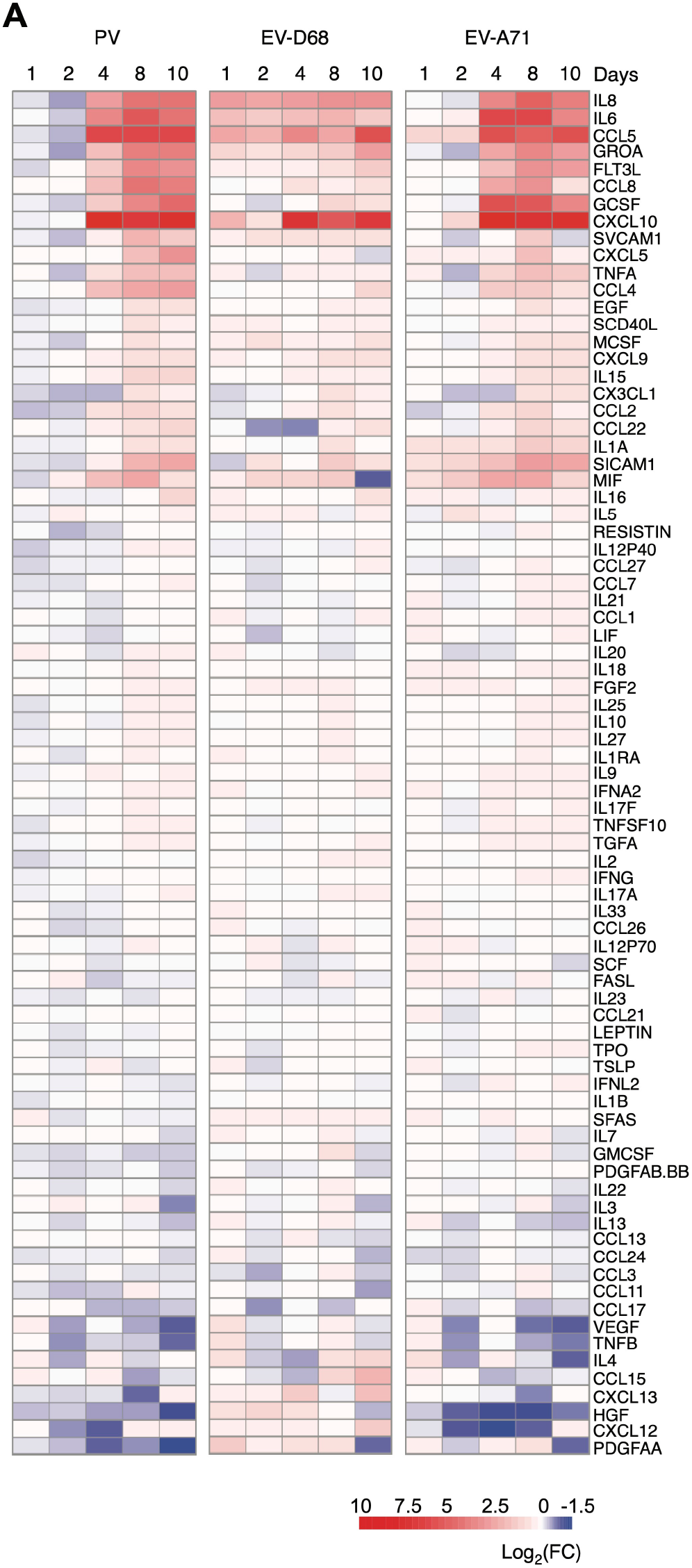
Cytokine profiling of EV infected hSpO, Related to Figure 2. **(A)** Luminex-based multianalyte profiling of 80 cytokines and chemokines of supernatants from EV infected hSpO at indicated days post infection. Cytokines are graphed as log2 fold change of mean fluorescent intensity (MFI) over uninfected controls at matched timepoint. MFI is the average of two technical replicates. Red denotes increased cytokines in comparison to uninfected, blue denotes decreased cytokines compared to uninfected and white denotes little to no change.

**Supplemental Figure 2:**
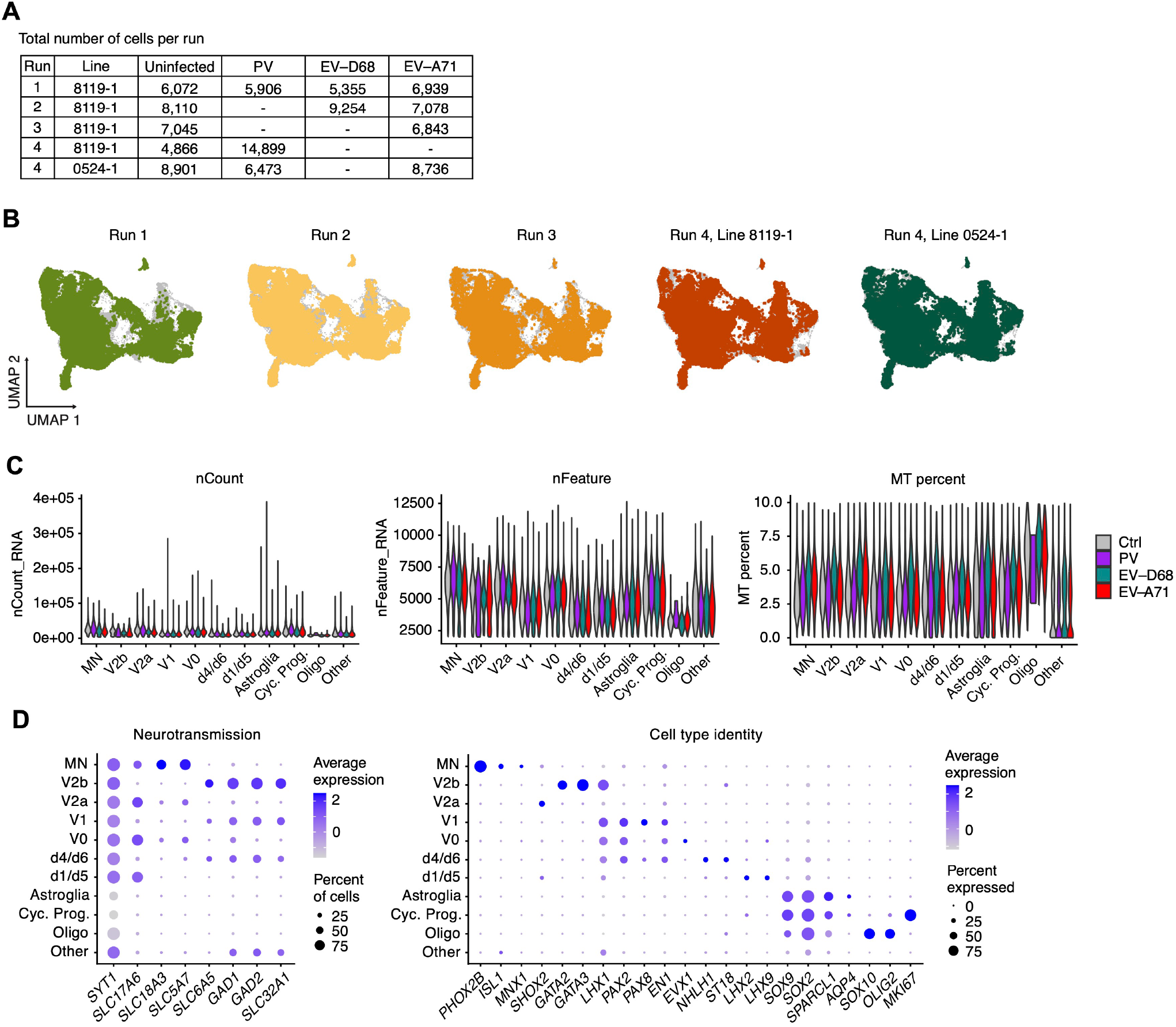
Quality metrics of single cell sequencing runs, Related to Figure 3. **(A)** hiPS cell lines used in each single cell run. Total number of cells from each enterovirus infected sample or uninfected sample per single cell run. **(B)** UMAP plots showing cells separately colored by the single cell run they were derived from. **(C)** Violin plots of total RNA count, gene count and mitochondrial gene content per each cell type cluster across uninfected and infected samples. **(D)** Dot plots showing the expression of selected neurotransmitter and cell-specific identity related genes. The size of the circle represents the percent of cells expressing each gene per cluster.

**Supplemental Figure 3:**
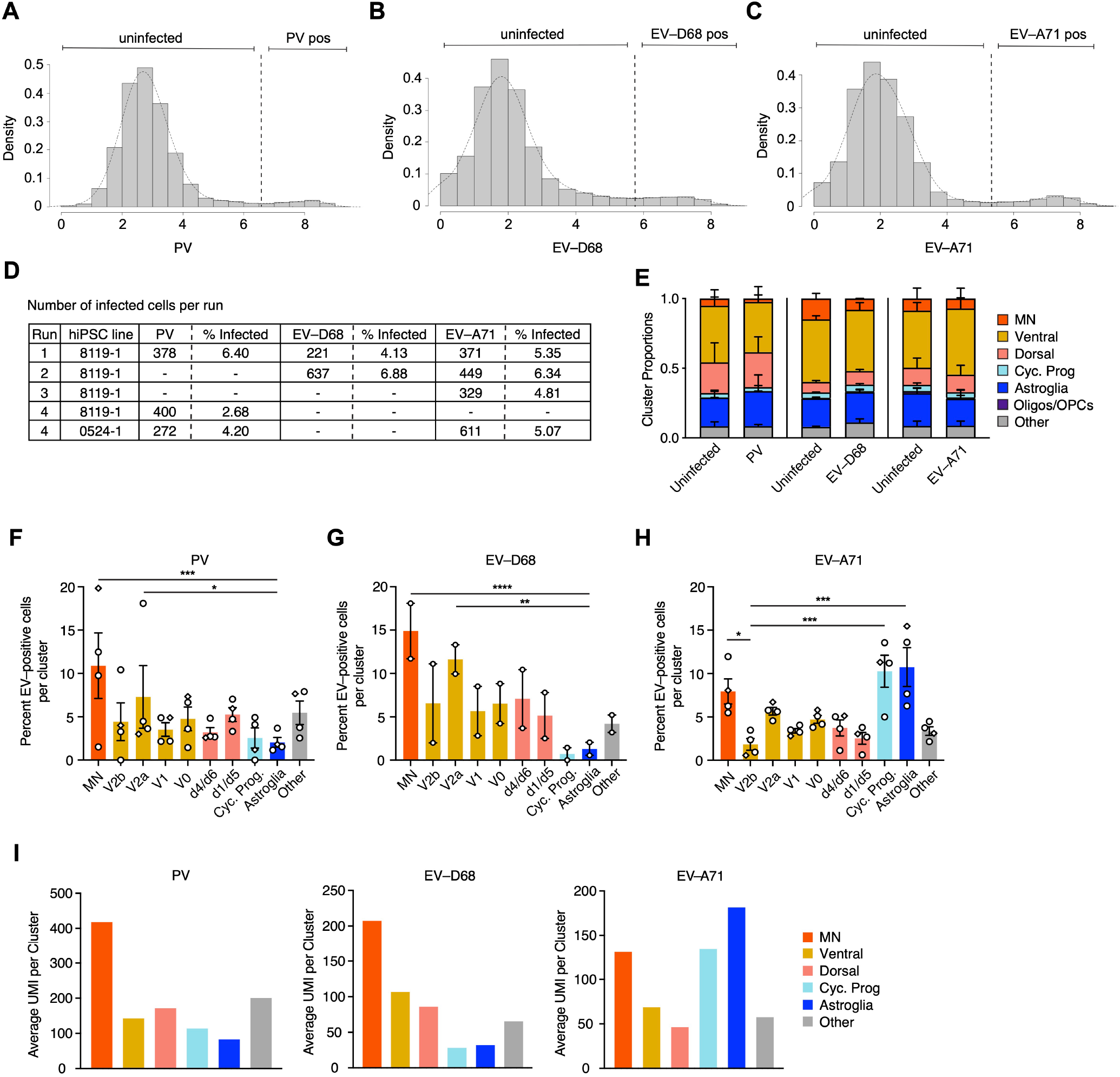
Identification of enterovirus infected cells by single cell sequencing, Related to Figure 3. **(A-C)** Histograms of virus expression for PV, EV-D68 and EV-A71 samples in scRNA-seq experiment. Histograms show bimodal distribution of virus expression. Dotted line indicates set threshold to discriminate between infected cells supporting active virus replication versus those containing viral RNA due to other processes (uninfected cells). **(D)** Distribution of the number of infected cells across each single cell run. **(E)** Graphs showing the percentage of cells from each sample belonging to each cluster. Each virus infected condition is matched to the uninfected samples from the same single cell runs. Datasets represent mean ± SD. **(F-H)** Percentage of infected cells per cell type in PV, EV-D68 and EV-A71 samples as graphed in Figure 3 separated by individual cell cluster. Datasets represent mean ± SEM. P-values were determined by two-way ANOVA with Benjamini and Hochberg correction. *P<0.05, **P<0.01, ***P<0.001, ****P<0.0001. **(I)** Average of the raw UMI counts of indicated enterovirus per cell type across all cells in PV, EV-D68 and EV-A71 infected samples.

**Supplemental Figure 4:**
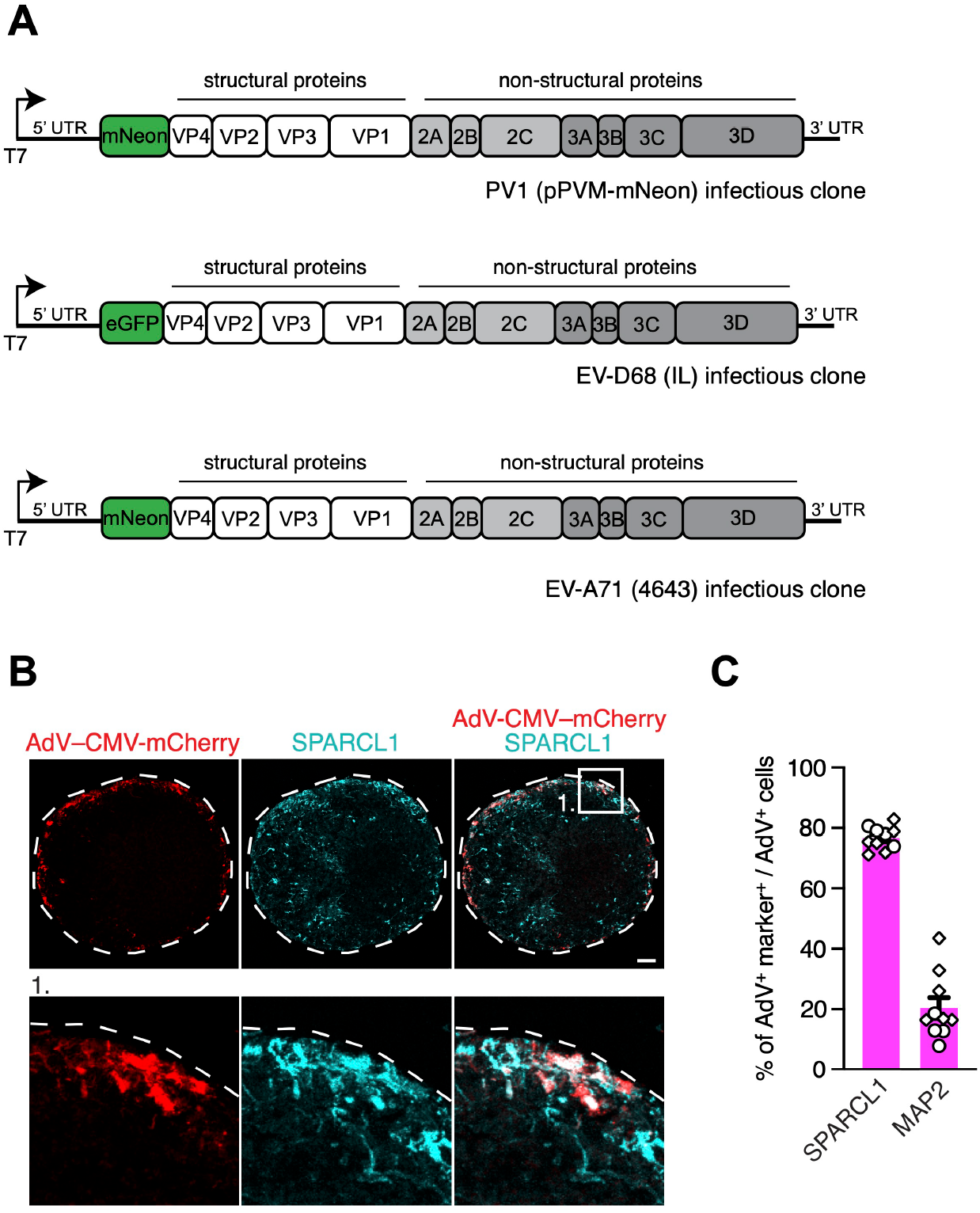
Validation of live imaging with reporter viruses, Related to Figure 4. **(A)** Schematic depiction of the PV1 infectious clone expressing mNeon with a GPI anchor tag, the EV-D68 (IL) infectious clone expressing eGFP, and the EV-A71 (4643) infectious clone expressing mNeon. T7, T7 promoter, UTR, untranslated region. **(B)** Representative SPARCL1 immunostainings of hSpO infected with AdV-CMV-mCherry. Scale bar, 100 μm. **(C)** Quantifications of AdV-CMV-mCherry co-localization with the neuronal maker MAP2 or glial marker SPARCL1. n= 10 hSpO from 2 hiPS cell lines from 1 differentiation. Datasets represent mean ± SEM.

**Supplemental Figure 5:**
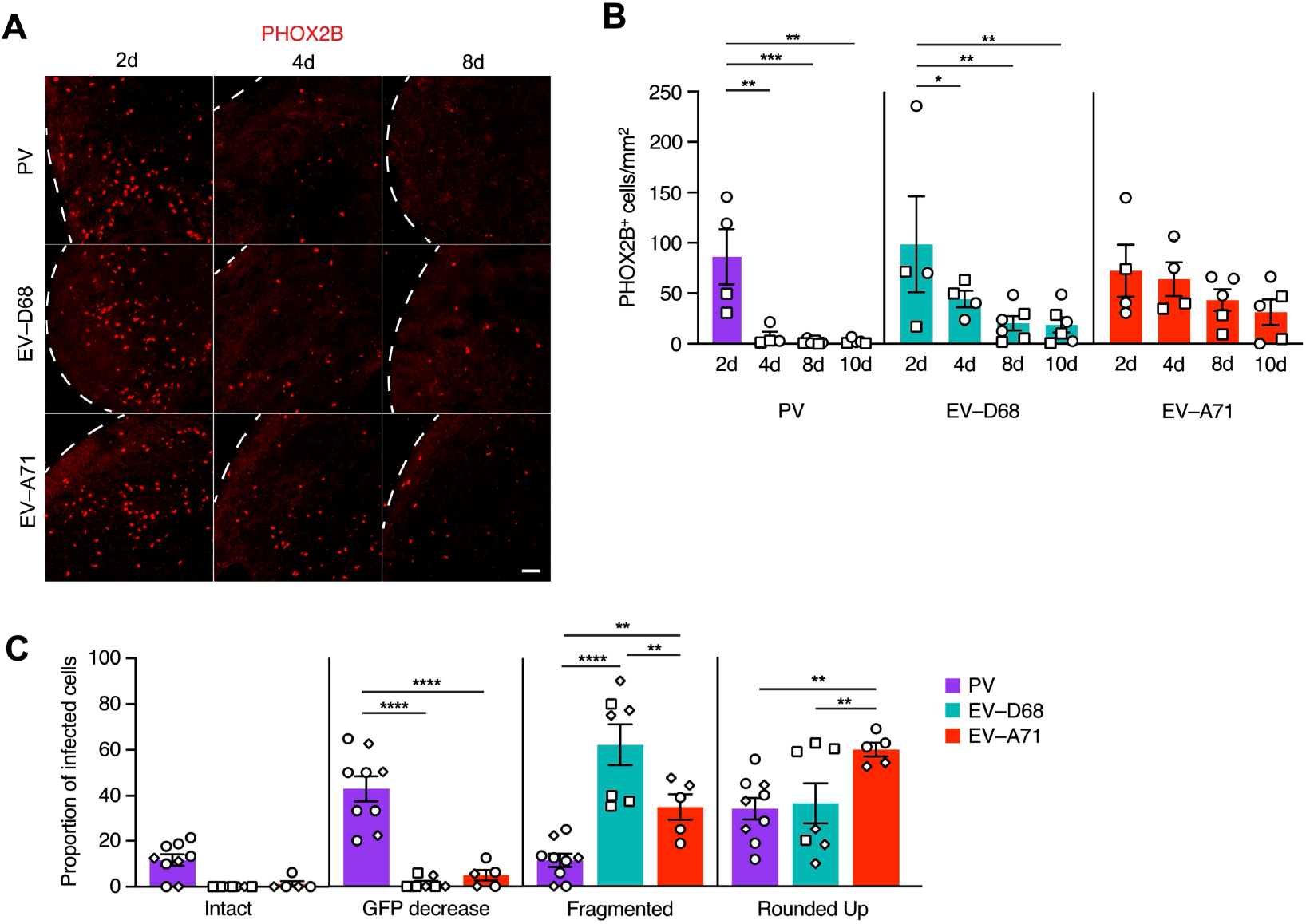
Cell damage in EV-infected hSpO, Related to Figure 5. **(A)** Representative PHOX2B immunostainings of hSpO infected with PV, EV-D68, or EV-A71 at indicated days post infection. Scale bar, 50 μm. **(B)** Number of PHOX2B positive cells per hSpO area for PV, EV-D68, or EV-A71 infected hSpO. n= 19 PV infected hSpO, n= 20 EV-D68 infected hSpO, n= 18 EV-A71 infected hSpO from 2 hiPS cell lines from 2 differentiations. Values represent mean ± SEM. P-values were determined by two-way ANOVA (adjusted with Benjamini–Hochberg). *P < 0.05, **P < 0.01, ****P < 0.0001. **(C)** Damage bins as related to Figure 5B, with individual values graphed. n= 9 PV infected hSpO from 2 hiPS cell lines from 1 differentiation, n= 7 EV-D68 infected hSpO from 2 hiPS cell lines from 2 differentiations, n= 5 EV-A71 infected hSpO from 2 hiPS cell lines from 2 differentiations. Datasets represent mean ± SEM. P-values were determined by two-way ANOVA (adjusted with Benjamini–Hochberg). **P < 0.01, ****P < 0.0001.

**Supplemental Video 1**. Spontaneous muscle contractions in uninfected hSpO-hSkM.

**Supplemental Video 2**. Spontaneous muscle contractions in PV infected hSpO-hSkM.

**Supplemental Video 3**. Spontaneous muscle contractions in EV-D68 infected hSpO-hSkM.

**Supplemental Video 4**. Spontaneous muscle contractions in EV-A71 infected hSpO-hSkM.

